# Spontaneous Emergence and Turnover of the Sex-Determining Locus

**DOI:** 10.64898/2026.01.26.701609

**Authors:** Takahiro Sakamoto, Hideki Innan

## Abstract

Anisogamy is nearly universal across multicellular organisms, yet how two discrete sexes arise in genomes with many recombining autosomal loci remains unresolved. Classic models show that disruptive selection on gamete size can drive anisogamy, but their single-locus assumptions conflict with empirical evidence that sexual differentiation typically involves numerous autosomal genes whose linkage is continuously broken by recombination. Here we develop an evolutionary model of gene regulatory networks (GRNs) to show that disruptive selection favoring complementary mating phenotypes can spontaneously generate a bistable developmental architecture producing two stable sexual phenotypes, together with a master sex-determining locus that biases development toward one or the other. Once such a bistable GRN evolves, the identity of the master regulator becomes flexible, enabling turnover of sex-determining loci without disrupting sexual differentiation. Our results provide a general mechanism for the origin, robustness, and evolutionary lability of sex-determination systems, linking classical gamete-level selection with the emergence of developmental bistability through the evolution of GRNs.

---

**A**nisogamy, the production of large ova and small sperm, predominates in multicellular species. It is a fundamental cause of sex differences^1^, and how it evolves from isogamy remains a central question in evolutionary biology. Disruptive selection favoring both large and small gametes is a key condition for its evolution. Classic theory shows that a size-number tradeoff in gamete production, combined with the survival advantage of larger zygotes, generates disruptive selection that drives anisogamy^2^. Subsequent work has expanded this framework^3–11^, and identified additional drivers, including size-dependent motility^12–14^ and cytoplasmic conflict^15–17^ (see reviews^18–20^). These studies collectively suggest that disruptive selection can arise under diverse conditions and may explain the widespread occurrence of anisogamy. However, most models rely on single-locus or asexual assumptions, leaving their applicability to multi-locus systems with recombination unclear.

Extending anisogamy theory to multi-locus settings is essential because sexual differentiation depends on many autosomal genes. Numerous studies show that multiple autosomal loci contribute to sexual differentiation through sex-biased expression, which underpins the pronounced morphological and functional divergence between the sexes^21–23^. However, extending single-locus models to multi-locus contexts is far from straightforward, because recombination disrupts the linkage of sex-specific alleles, generating maladaptive intermediate phenotypes and hindering the evolution of discrete sexes. One possible solution is the clustering of sex-related genes into tightly linked regions, such as sex chromosomes or supergenes^4,24^. Yet this explanation contrasts with empirical evidence that sexual differentiation commonly involves many autosomal genes dispersed across the genome, highlighting the need for theory that accommodates multi-locus architectures with recombination.

We hypothesized that this paradox can be resolved by the developmental dynamics encoded in a gene regulatory network (GRN). Our previous work showed that GRN evolution can generate discrete phenotypic plasticity even under recombination^25^. Here, building on this framework^25–27^, we ask how disruptive selection favoring complementary mating pairs can drive the emergence of two discrete sexes from a monomorphic population. In our model, sexual differentiation, the emergence of two stable gene expression states corresponding to alternative sexes, evolves when the system acquires a bistable GRN with alternative developmental equilibria and a master sex-determining locus that directs development toward one equilibrium or the other. This mechanism aligns with empirical systems in which mutually antagonistic feedback loops underlie sexual differentiation^28,29^. In this study, we investigate how such a bistable GRN and a master sex-determining locus can arise spontaneously through evolution.

Our model points to an inherent flexibility in the identity of the master regulator of sexual fate, mirroring the diversity of master sex-determining factors from single locus to polygenic control^29^ and the recurrent turnover of sex-determination systems^28,30^. Although bistable GRNs are thought to underlie such flexibility^29^, previous theories^31–33^ presupposed master loci and downstream cascades, and lacked a formal mechanism for how flexibility could arise. In our framework, however, sexual differentiation is driven by the feedback architecture of the GRN rather than by any single locus, allowing multiple genes to adopt the role of master regulator without disrupting bistability. This mechanism explains observed turnovers in sex-determining loci^28,30^ and predicts that turnover can buffer the deleterious effects of sex-chromosome degeneration^34,35^ by replacing degenerated loci with new, benign regulators^36,37^. Together, these results identify feedback architecture as the key principle underlying the origin, robustness, and flexibility of sex-determining systems.

## Results

### Evolutionary model of gene regulatory network

Building on the idea that developmental dynamics governed by a GRN shape the evolution of anisogamy, we employ a simple GRN model, illustrated in Fig. 1 (see Methods). A population of *N* diploid individuals undergoes repeated cycles of development, mating, and inheritance. During development, gene interactions determine phenotypes; during mating, disruptive selection acts on these phenotypes; and through inheritance with mutation and recombination, GRNs evolve over generations.

**Fig. 1:**
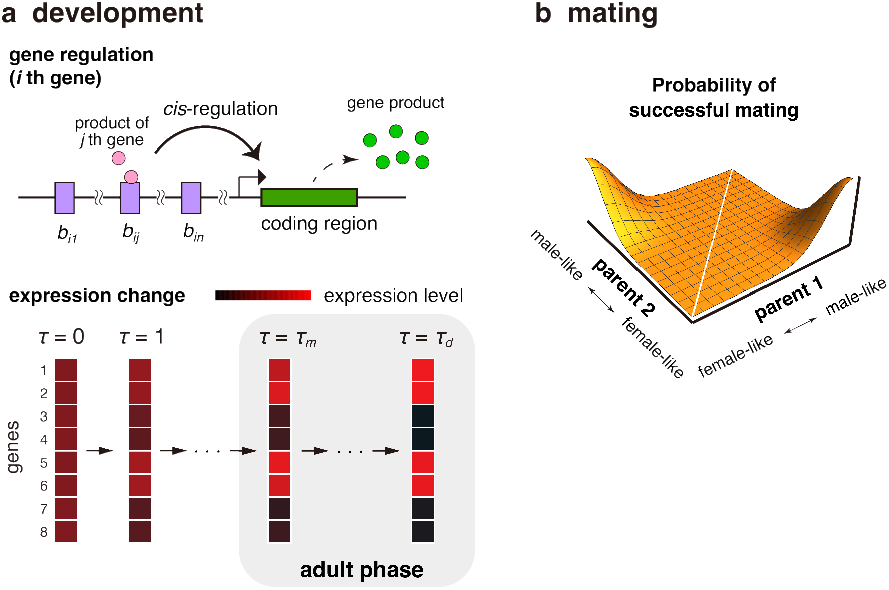
Overview of the model. Each generation consists of development and mating. (a) In the development event, each gene expresses under the control of the type-B loci (upper panel). Defined by the gene regulation, the expression levels change from birth (*τ* = 0) to death (*τ* = *τ*_*d*_) of each individual (lower panel). Expression levels during the adult phase (*τ*_*m*_ ≤ *τ* ≤ *τ*_*d*_) represent the individual’s phenotype and are subject to selection in the mating event. (b) In the mating event, disruptive selection is assumed such that mating is more likely to be successful when one individual has a male-like phenotype while the other has a female-like phenotype.

Each individual carries *n* autosomal coding genes whose expression levels define the phenotype (Fig. 1a). Gene expression is controlled by regulatory interactions among genes: the initial expression of gene *i* is set by a type-G locus (*g*_*i*_), and its response to gene *j* is determined by type-B loci (*b*_*ij*_; *j* ∈ {1, …, *n}*). These allelic values jointly specify the developmental dynamics (equation (1) in Methods). Development proceeds from the embryonic stage (*τ* = 0) to adulthood *τ* = *τ*_*m*_ and continues until death *τ* = *τ*_*d*_. Adult phenotypes are subject to disruptive selection associated with reproductive roles. We focus on evolutionary changes in gene expression levels and ignore the evolution in coding sequences.

During mating, selection favors pairs in which one individual performs well in the male function and the other in the female function (Fig. 1b). The optimal expression profiles for the two roles are given by vectors 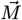 and 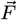. An individual’s fitness as a male (*W*_*m*_) or female (*W*_*f*_) depends on the similarity between its phenotype and the corresponding optimum (ranges from 0 and 1; see Methods). Mating pairs form randomly, but their probability of reproducing successfully increases when one partner has high *W*_*m*_ and the other high *W*_*f*_. Each successful pair produces one offspring, and reproduction continues until the next generation reaches population size *N*.

Inheritance follows free recombination among all *g*_*i*_ and *b*_*ij*_ loci. Type-G and type-B loci mutate at rates *µ*_*g*_ and *µ*_*b*_ per locus, and mutations are introduced into offspring.

Using simulations based on this model, we tested whether GRN evolution in autosomal genes under free recombination can generate dimorphic sexes and ultimately produce anisogamy. Each simulation began with an isogamous, homogeneous population in which all individuals shared the same genotype and had *W*_*m*_, *W*_*f*_ ≪ 1 (Methods). We considered sexual dimorphism to have evolved when all individuals adopted either male or female roles with no intermediates, i.e., when max(*W*_*f*_, *W*_*m*_) > 0.95 for all individuals. In this study, we used *n* = 8 autosomal genes and a constant population size of *N* = 1,000.

### Establishment of sexual dimorphism

We first examine whether differentiated male and female phenotypes evolve in our model. To do so, we ran simulations across a range of parameter values, varying one focal parameter (mutation rate, selection strength, or optimal expression profiles) while keeping the others at default values (Table S1). For each parameter set, we performed 1,000 simulation replicates. Each simulation was run for up to 5×10^6^ generations, or until dimorphic sexes were established, defined as max(*W*_*f*_, *W*_*m*_) > 0.95 for all individuals. We then recorded, for each parameter set, the proportion of runs in which anisogamy emerged (*P*_*a*_) and the distribution of waiting times (*T*).

Across all parameter sets (Table S1), two distinct sexes emerged in most simulation runs (*P*_*a*_ ≈ 1; Fig. S1). This highlights the capacity of the GRNs to generate sexual dimorphism spontaneously, even when the underlying genes are unlinked. Among the examined parameters, *µ*_*b*_ had a noticeable effect: when *µ*_*b*_ was small, *P*_*a*_ declined slightly (∼95%) because the regulatory network can evolve too slowly for dimorphism to arise within 5×10^6^ generations. Consistently, the waiting time *T* increased as *µ*_*b*_ decreased. In contrast, selection strength and optimal expression patterns had little effect on either *P*_*a*_ or *T*, and *µ*_*g*_ showed no notable effect.

### Evolutionary trajectory toward sexual dimorphism

To characterize how sexual dimorphism arises and is maintained within simulations, we examined the evolutionary dynamics across simulation replicates and found highly consistent patterns. We therefore focus on a representative run under the default parameter set, which illustrates a typical evolutionary trajectory. Fig. 2 shows the evolutionary dynamics in a representative run in which the first four genes share the same optimum, while the remaining four are under sex-specific divergent selection (Fig. 2a, see also Table S1). The frequencies of male-like and female-like individuals—defined by *W*_*m*_ > 0.95 and *W*_*f*_ > 0.95, respectively—are shown in Fig. 2b. After the two sexes emerge at *T* ≈ 165,000, the population maintains a stable 1:1 sex ratio. We examine how development proceeds within individuals and how it evolves over time. Fig. 2c shows developmental trajectories of gene expression at five evolutionary time points (indicated by arrows in Fig. 2b). To generate this plot, we recorded genotypes every 10,000 generations, computed phenotypes along development, and summarized the resulting phenotypic distribution using principal component analysis (PCA). The first two principal components captured ∼99% of total phenotypic variance. PC1 was almost perfectly aligned with 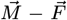 and therefore captured sexual differentiation, whereas PC2 mainly reflected coordinated expression of the first four genes, a shared-optimum trait (e.g., a general growth-related trait). Developmental steps (*τ*) are shown by color shading in Fig. 2c. Together, these trajectories reveal the gradual emergence of sexual dimorphism along PC1: phenotypes remain largely overlapping at *t* = 10,000, show clear separation by *t* = 50,000, become strongly polarized by *t* = 500,000, and are stably maintained once the two sexes are fully established (e.g., at *t* = 2,000,000).

**Fig. 2:**
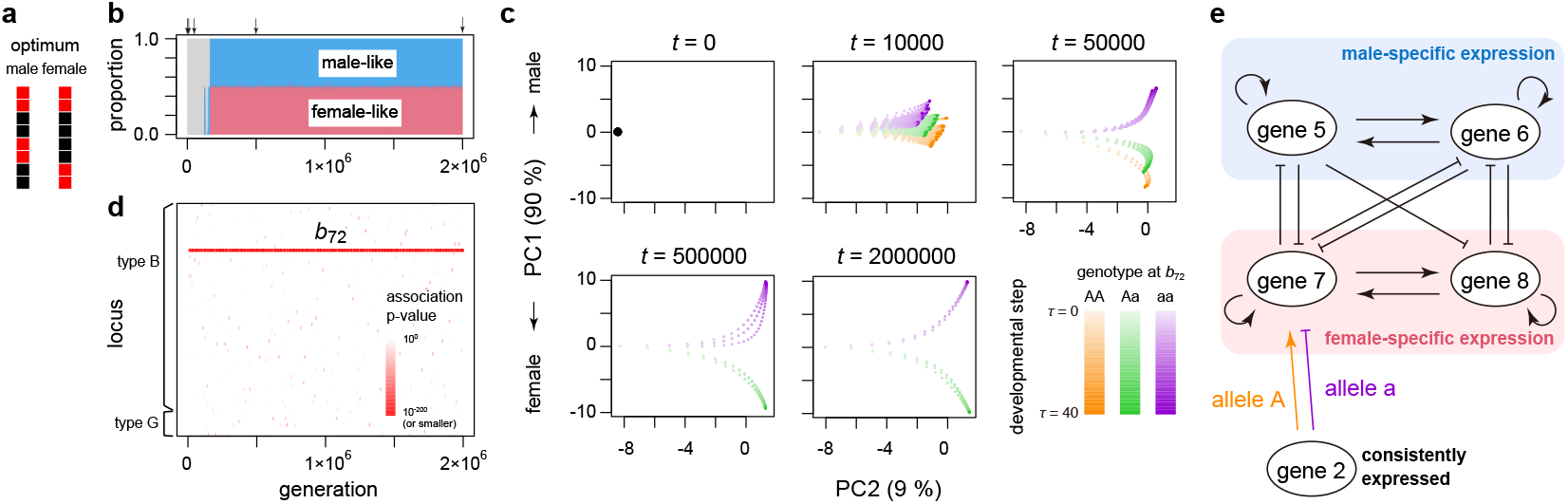
Typical evolutionary dynamics of the emergence of two sexes. The result of a representative simulation run with the default parameter set is shown (see Table S1). (a) The optimal expression pattern of each sex. (b) The population frequencies of male-like and female-like individuals (in blue and red, respectively). The gray region represents the proportion of individuals with intermediate phenotypes. Arrows on the top shows five time points focused in panel c. (c) Phenotypic changes during development of all individuals surviving at the five time points. The shade of circles represent the developmental step (*τ*). Colors show the genotype at *b*_72_, which is the major sex-determining locus in this simulation run. (d) Association between the sex-specific phenotype and genotype at each locus. The p-values from Kendall’s *τ* -*b* rank correlation test were computed for all loci (*n* type-G loci and *n* × *n* type-B loci (*b*_*ij*_ with *i, j* = 1, …, *n*) from bottom to top). (e) Schematic view of gene regulation underlying sexual differentiation. The graph is drawn based on the genotype at *t* = 2,000,000, where regulatory relationships with |*b*_*ij*_ | > 0.05 are shown. The relationships unrelated to the sexual dimorphism are ignored here. For the full regulatory relationship, see Fig. S2.

Interestingly, variation in developmental trajectories among individuals is well explained by focusing on a specific locus (*b*_72_ in this run). Alleles A and a denote variants with higher and lower regulatory values, respectively. In Fig. 2c, individuals are colored by their genotypes at *b*_72_. By *t* = 10,000, allele A already biases development toward female-like phenotypes, though substantial within-genotype variation indicates contributions from other loci. By *t* = 50,000, sexual differentiation has advanced, and *b*_72_ becomes the primary determinant, as shown by reduced variation within genotypes. At this stage, genotype AA becomes rare because matings between individuals carrying allele A—who tend to develop female-like phenotypes—are largely unsuccessful. By *t* = 500,000, a ZW-like system is established, with females as heterozygotes (Aa) and males as homozygotes (aa), and this pattern remains stable through *t* = 2,000,000.

Indeed, *b*_72_ acts as the sex-determining locus that emerged in this run. Such loci can be readily identified from adult phenotypes as those showing strong associations with sexspecific traits (see Methods). Fig. 2d shows the association strengths for all *n* type-G loci and *n*^2^ type-B loci. *b*_72_ begins to exhibit a pronounced association early in the simulation.

The role of *b*_72_ in the GRN is straightforward (Fig. 2e). Focusing on genes with sex-specific optima (genes 5–8), their regulatory interactions (*b*_*ij*_, *i, j* ∈ {5, 6, 7, 8}) form a mutually inhibitory structure: genes 5 and 6 activate themselves and repress genes 7 and 8, and vice versa. This architecture makes the expression of the two gene pairs (5 and 6 vs. 7 and 8) mutually exclusive, producing bistable developmental dynamics corresponding to male and female optima. Within this network, *b*_72_ breaks the symmetry: its genotype (alleles A or a in Fig. 2e) determines whether gene 2 enhances or represses gene 7, thereby selecting the developmental equilibrium. Importantly, gene 2 has the same high-expression optimum in both sexes. Male and female genotypes differ almost solely at *b*_72_ (Fig. S2), indicating that *b*_72_ functions as the primary sex-determining switch.

### Frequency distribution of sex-determining loci in the regulatory network

Given the results above, we hypothesized that sex-determining loci are likely to occur at loci *b*_*ij*_ where *i* is one of the genes with sex-specific optima (5–8) and *j* is a constitutively expressed gene (1 or 2), allowing the latter to act as a reliable trigger. Consistent with this expectation, loci with *i* ∈ {5, 6, 7, 8} and *j* ∈ {1, 2} served as sex-determining loci in most simulation runs under the default parameter set (Fig. 3). This distribution was largely unchanged across different mutation rates and selection strengths (Fig. S3), indicating that the prediction is robust.

**Fig. 3:**
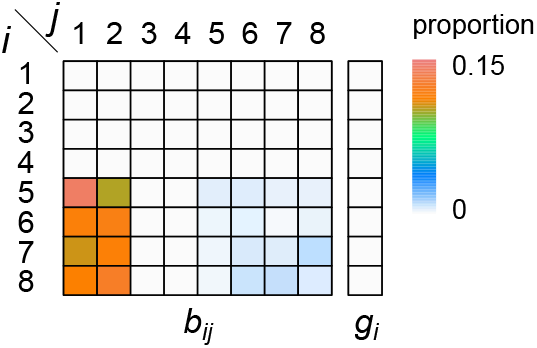
Frequency distribution of sex-determining loci. The frequency distribution of sex-determining loci at generation *t* = *T* is obtained from 998 runs with the default parameter set, in which the two sexes have established within 5 × 10^6^ generations. For the results with other parameters, see Fig. S3.

These findings imply that constitutively expressed genes often play a central role in anisogamy evolution; however, this role is not essential. When we assumed that all genes experience sex-specific selection, and thus no constitutively expressed regulators exist, sex-determining loci still arose. Under these conditions, all type-B locus were approximately equally likely to serve as a regulator (Fig. S3).

### Turnover events of sex-determining locus

We next analyzed how sex-determining loci changed over time in the simulations. To do so, we extended the simulations in which anisogamy had evolved, continuing each run for additional 5×10^6^ generations after dimorphic sexes were first established (i.e., until *t* = *T* + 5 × 10^6^).

With the default parameter set, a single locus typically works as a sex-determining locus, and polygenic sex determination appeared only transiently. Most runs maintained the same locus across the entire simulation (Fig. 2c), but some exhibited turnover events (Fig. S4). These turnovers generally preserved the 1: 1 sex ratio and often seemed driven by random genetic drift. Transitions occurred either without a change in the heterogametic sex (Fig. S4a) or with a switch between XY and ZW systems (Fig. S4b); some replicates showed multiple turnovers in a short interval (Fig. S4c).

Fig. 4 shows how a turnover of sex-determining loci occurred without disrupting the 1: 1 sex ratio in a representative run. In this run, the sex-determining locus shifts from *b*_51_ to *b*_71_ (Fig. S4a). Denote the alleles at these loci by A/a and B/b, with capital letters representing larger genotypic values (i.e., leading to higher expression). Here, alleles a and B work as feminizing alleles, and the turnover does not change the heterogametic sex (ZW system). The total frequency of female-producing genotypes (Aabb and AABb) remained close to 0.5 throughout the turnover (Fig. 4a). Comparisons of regulatory interactions before and after the turnover (Fig. 4b,c) show that the bistable module among genes 5–8 remains unchanged; only the upstream trigger differs. In the ancestral system, feminization was achieved by repression of gene 5, whereas in the derived system it occurs through activation of gene 7. This demonstrates that multiple loci (most likely *b*_*ij*_ for *i* ∈ {5, 6, 7, 8} and *j* ∈ {1, 2} in this setting) can potentially act as sex-determining loci while the underlying bistable differentiation mechanism remains conserved.

**Fig. 4:**
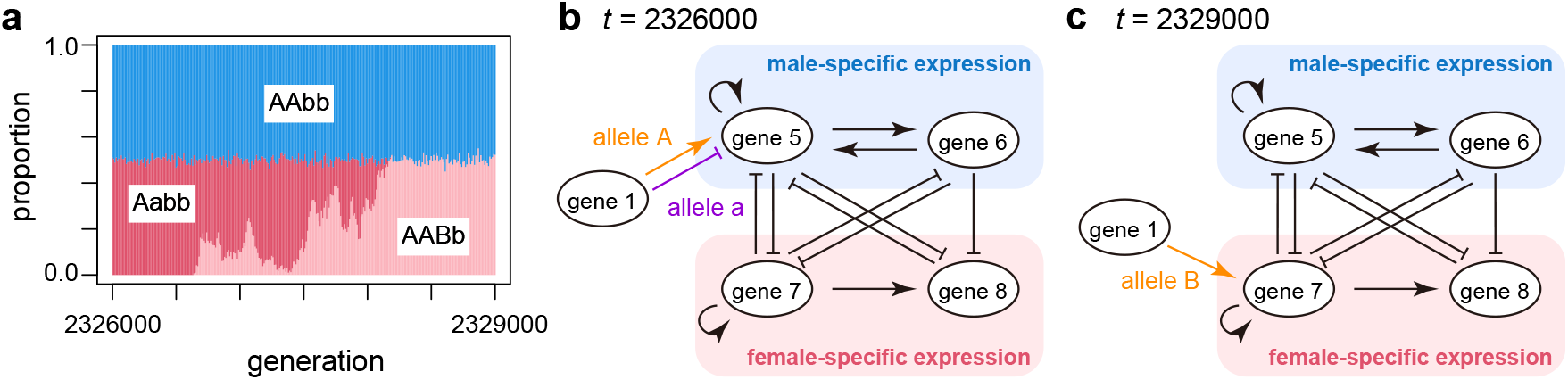
Turnover of a sex-determining locus. As an example, the simulation run shown in Fig. S4a is illustrated. (a) The relative frequencies of three genotypes (AAbb, Aabb, and AABb) during the turnover. Other genotypes are very rare and colored in gray. (b, c) Gene regulatory relationships involved in the sexual differentiation before (b) and after the turnover (c). Gene regulations with |*b*_*ij*_ | > 0.05 are shown for *i, j* ∈ {5, 6, 7, 8}.

Turnover events can occasionally involve a change in the heterogametic sex. We observed dynamics qualitatively similar to those observed in cases without changing heterogametic sex, and an example is shown in Fig. S5. In this case, the sex-determining role shifts from *b*_52_ to *b*_51_, accompanied by a transition from an XY to a ZW system (Fig. S4b). In the ancestral system, allele A is masculinizing, and a new allele B arises that is also masculinizing in homozygotes. This turnover involves complex frequency dynamics across the six relevant genotypes (Fig. S5a), yet the proportions of males and females remain close to 0.5. As before, the regulatory interactions among genes 5–8 show no qualitative changes; only the upstream trigger of sexual determination differs (Fig. S5b, c).

### Rate of turnovers of sex-determining locus

To quantify the overall patterns, we analyzed the dynamics across all replicates and calculated the proportion of runs exhibiting a turnover of the sex-determining locus (*P*_*t*_; Fig. S1c). Under the default parameter set, 157 out of 998 replicates showed at least one turnovers, producing 814 turnover events in total. Among these events, 726 retained the heterogametic sex and 88 altered it.

We next examined how each parameter in Table S1 affects the turnover frequency of the sex-determining locus. Among them, the mutation rate at type-B loci (*µ*_*b*_) had the strongest effect: the turnover proportion (*P*_*t*_) increased with higher *µ*_*b*_, likely because it raises the supply of potential sex-determining alleles. Selection strength and expression optima had more moderate effects. Stronger selection reduced turnover, as a new sex-determining locus must be sufficiently functional to replace the existing one. Similarly, *P*_*t*_ decreased when sex-specific expression applied to all genes, probably because anisogamy under this condition requires more finely tuned master regulators, increasing selection against alternatives. The mutation rate at type-G loci had little effect. Turnovers that altered the heterogametic sex were generally less common than those that did not.

### Recovery from degenerated sex chromosomes

So far, our results show that, through the evolution of a bistable GRN, a single locus typically evolves to function as the sex-determining locus. This locus usually retains two alleles for prolonged periods unless a turnover occurs. Consequently, the chromosomes harboring this locus can be regarded as proto-sex chromosomes. It is well established that, once sex chromosomes emerge, the Y (or W) chromosome accumulates deleterious mutations after recombination suppression. We here propose that GRN flexibility provides long-term protection against such degeneration. Turnovers of sex-determining loci may enable populations to recover from, or avoid, Y/W degeneration^31,36,37^ (Fig. 5a).

**Fig. 5:**
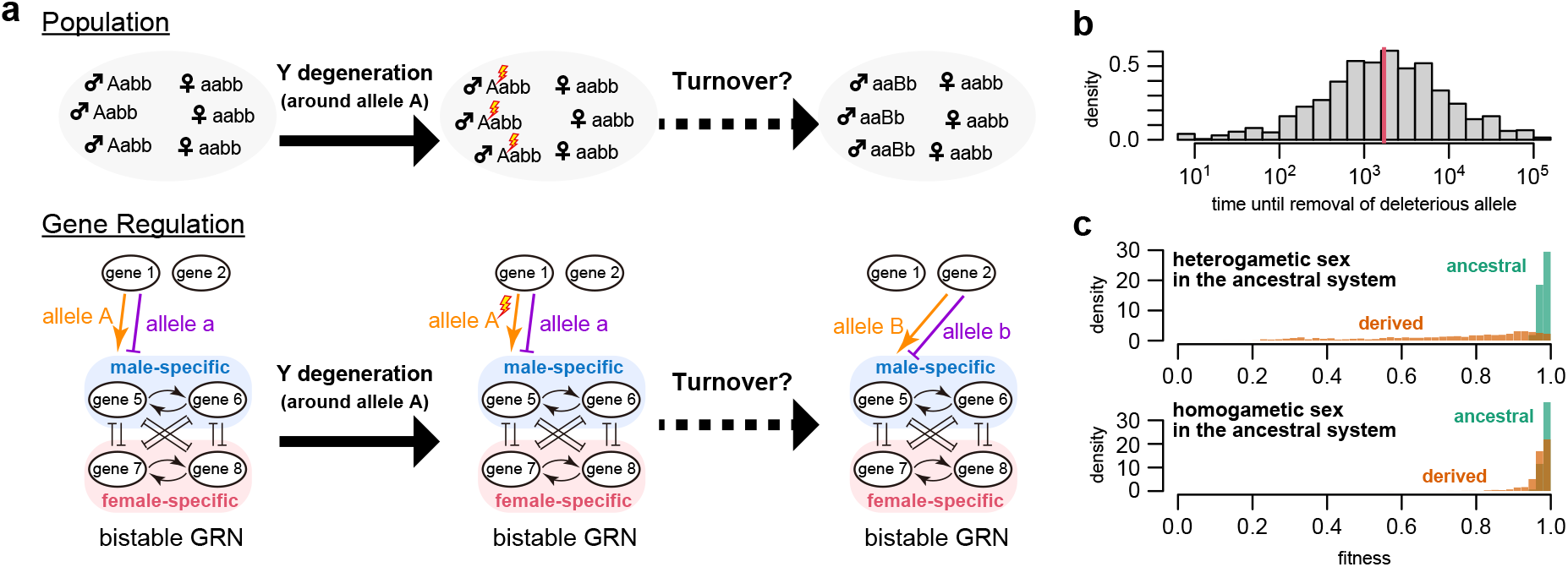
Recovery from degenerated sex chromosomes. (a) Schematic view of a recovery process from degeneration through a turnover of sex-determining allele. In the ancestral system, a regulatory locus, *b*_51_, determines sex: allele A enhances the expression of gene 5 whereas allele a represses it (left panel). Here, the chromosomes carrying allele A and a are regarded as Y and X, respectively. Once recombination is suppressed around *b*_51_, deleterious mutations can rapidly accumulate on the Y chromosome (middle panel). Because these mutations are tightly linked to the sex-determining allele A, purging them without removing allele A is difficult. However, the emergence of a new sex-determining locus (e.g., *b*_52_) could allow the removal of both the deleterious mutation and its associated sex-determining allele without a large fitness cost (right panel). (b) Distribution of the number of generations required to remove the deleterious allele after a highly deleterious allele is introduced in complete linkage with the sex-determining allele on Y or W chromosomes. The distribution is based on 1,000 simulation runs where the two sexes are established within 5 × 10^6^ generations. The red vertical line shows the median. (c) Comparison of fitness just before and after the removal of the deleterious allele. The mean fitness of the ancestrally heterogametic and homogametic sexes is shown in each panel.

To test this hypothesis, we designed simulations in which a strongly deleterious mutation was forcibly fixed near the dominant sex-determining allele on the Y or W chromosome (allele A in Fig. 5a), while the recessive allele on the X or Z chromosome (allele a) was left intact (see Methods), thereby mimicking degeneration. We then examined whether the system could purge this mutation by recruiting a new sex-determining locus. Although somewhat artificial, this setup provides a useful framework for evaluating whether the system can replace a degraded sex-determining locus.

Under the default parameter set, the deleterious allele was removed in all 1,000 replicates. The process involved two phases: (i) the appearance of a new sex-determining allele and (ii) its subsequent spread, during which the ancestral allele carrying the deleterious mutation was lost. Phase (i) required a long waiting time, whereas once a new allele arose, it spread rapidly and removed the old one (Fig. S6a). The entire elimination process generally completed relatively quickly, with a median of ≈ 1,700 generations (i.e., on the order of *N*) though waiting times varied widely due to stochasticity (Fig. 5b, Fig. S6a). Although rare, elimination occurred almost instantaneously in some cases, especially when a new sex-determining allele was already present as standing variation. In most replicates (751/1,000), the new sex-determining allele arose at a different locus, resulting in a turnover.

After removal of the deleterious allele, the sex-determining system was often suboptimal, showing reduced fitness in both sexes (*W*_*m*_ and *W*_*f*_; Fig. 5c), particularly in the ancestrally heterogametic sex. This occurred because the ancestral system had deteriorated to the point where even a suboptimal alternative was selectively favored. The GRN then adapted to restore the original fitness level. Because the bistable regulatory architecture was already established, this recovery was much faster than the initial evolution of dimorphism from an isogametic state (Fig. S6b). In summary, recovery proceeded in two steps: a rapid turnover to a mutation-free but suboptimal system, followed by gradual refinement. This two-step pattern was consistent across parameter regimes (Fig. S7).

## Discussion

Sexual differentiation involves complex interactions among many genes across the genome, including those on autosomes. We show that a GRN can evolve to coordinate unlinked genes into two discrete, stable sexual phenotypes when disruptive selection acts on phenotype. This framework provides a general mechanism for the origin of anisogamy and recapitulates key features of natural sex-determination systems^28^: (i) sex is typically controlled by a single master gene, (ii) the identity of this gene is evolutionarily flexible, allowing turnovers, and the downstream sexual differentiation mechanism remains conserved despite such turnovers.

In our model, mutually antagonistic feedback loops constitute the core mechanism that allows flexibility in sex-determining factors while maintaining robustness in downstream sexual differentiation. Within this architecture, early developmental differences in gene expression are amplified as development progresses, enabling alternative sexual phenotypes. This suggests that many genes influencing early expression have the potential to contribute to sex determination. Once a developmental fate is specified, the phenotype is stabilized through a positive feedback loop (Fig. 2e). This canalization prevents intermediate phenotypes even when new sex-determining alleles or loci arise, enabling turnover with minimal fitness costs^31,38^. Consistent with our result, such feedback architectures are common in empirical systems^28^, and our model helps explain their evolutionary origins.

Our model also predicts that this network architecture can prevent the evolutionary loss of sex that might otherwise occur due to highly degenerated sex chromosomes. Longterm retention of the same sex chromosomes is known to cause progressive degeneration as deleterious mutations accumulate^34,35,39^. Based on this, some researchers have even suggested a catastrophic scenario in which the loss of a sex-determining gene ultimately results in the disappearance of one sex — and possibly species extinction^40^. In contrast, our results show that the GRN can avoid such outcomes by recruiting new master regulators while purging deleterious mutations linked to the ancestral sex-determining gene^41,42^. This recovery proceeds through a rapid turnover to a new, initially suboptimal sex-determination system, followed by its gradual refinement. Although previous theories have suggested that selection may favor new systems after degeneration^31,36,37^, our model emphasizes the key role of transiently suboptimal sex-determining alleles. These states may act as evolutionary refugia, buffering the system from extinction risk and supporting long-term stability of sexual differentiation.

In this study, we focus on the role of GRNs in the early evolution of sexual differentiation systems, while recent theories emphasize gene regulation in the later evolution of sex chromosomes^43–45^. Those studies assume established dimorphic sexes and sex-specific regulatory programs, and investigate how genes on the sex chromosomes (i.e., genes linked to the sex-determining locus) acquire sex-specific expression. By contrast, our work addresses an earlier phase: how the regulatory architecture that supports dimorphic sexes can arise from an ancestral isogametic state. These frameworks are complementary, capturing different aspects of sex system evolution: our study examines the emergence and subsequent evolution of sex-determining loci, while these previous works describe the regulatory divergence in the sex-linked genomic regions. Integrating these perspectives in future research will be a promising direction toward a unified theory of sex chromosome evolution, spanning their origin, diversification, maintenance, and turnover.

## Methods

### Simulation model

We simulated a Wright–Fisher population consisting of *N* diploid individuals. In each generation, all individuals first undergo development, and then participate in mating, during which disruptive selection acts on phenotypes of mating pairs. The phenotype is defined by the expression levels of *n* autosomal coding genes, which change dynamically during development according to regulatory interactions among genes. The developmental model follows our previous formulation^25^. The gene regulatory network consists of two types of loci, type-G and type-B loci. Type-G loci determine the initial expression levels at the embryonic stage, whereas type-B loci define the regulatory interactions that govern subsequent expression dynamics. Let *x*_*i*_(*τ*) denote the total expression level of gene *i*, defined as the sum of the expression outputs from both copies. The initial expression level is

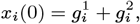

where 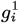 and 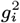 are genotypic values at the *i*th type-G locus inherited from the first and second parents, respectively. During development, expression levels follow

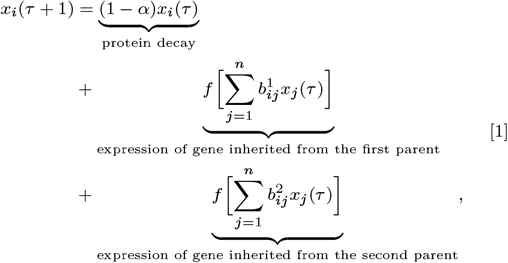

where 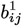 and 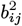 are the genotypic values at the type-B loci inherited from the first and second parents, respectively, and *α* is the per-step decay rate of the gene product. As equation (1) shows, we assume that type-B loci act in *cis*, influencing the expression of the gene copy inherited from the same parent but not the other. *f*(*x*) = (1+tanh(*x*))*/*2 is a sigmoid function that normalizes expression between 0 and 1. Under this formulation, the possible range of *x*_*i*_(*τ*) is 0 ≤ *x*_*i*_(*τ*) ≤ 2*/α*. We neglect stochastic expression noise during development. Together, type-G and type-B loci determine the developmental trajectory *x*(*τ*), which constitutes the phenotype in our model.

Reproduction occurs through mating between pairs of individuals. Any pair may mate, but the probability of successful mating depends on their phenotypes: it is maximized when one individual expresses the optimal male function and the other the optimal female function. To model this, we define optimal male and female phenotypes as vectors of ideal gene-expression levels. Let 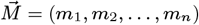 and 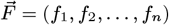 denote the male and female optima, respectively. An individual’s adult phenotype is given by 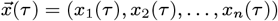, and its sex-specific performance is evaluated by comparing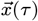 with the corresponding optimal vector. Let *W*_*i*_ denote performance as sex *i* (*i* ∈ {*m, f*}). Assuming that lifetime performance is the geometric mean over the adult developmental period, we define

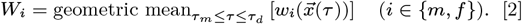

where the instantaneous performance is given by 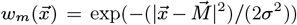 and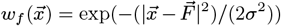. The probability of successful mating between individuals *i* and *j* is then proportional to

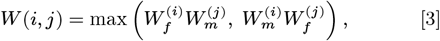

where 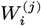 is the sex-*i* performance of individual *j*. In the Wright–Fisher reproduction step, pairs of individuals are sampled with replacement, and each pair produces one offspring with probability *W/* max(*W*).

After mating, mutations are introduced at both type-G and type-B loci. Each type-G locus mutates with probability *µ*_*g*_ per locus per generation, changing *g*_*i*_ by Δ*g* ∼ Uniform(−*γ*_*g*_, *γ*_*g*_). To ensure that allele expression levels remain within the biologically feasible range [0, 1*/α*], we impose boundaries: if *g*_*i*_ < 0, then *g*_*i*_ = 0; if *g*_*i*_ > 1*/α*, then *g*_*i*_ = 1*/α*. In practice, all *g*_*i*_ values remained far from these boundaries across all simulations, and thus this correction had no effect on the results. These parameters are constant across all *n* type-G genes. Mutation at type-B loci occurs independently: each *b*_*ij*_ mutates with probability *µ*_*b*_ per generation, with a mutational effect Δ*b*_*ij*_ ∼ Uniform(−*γ*_*b*_, *γ*_*b*_). Again, these parameters are uniform across all type-B loci. Free recombination is assumed between all pairs of *g*_*i*_ and *b*_*ij*_.

Each simulation run begins with a homogeneous population, where all individuals have the genotype: *g*_*i*_ = 1*/*(2*α*) and *b*_*ij*_ = 0 for all *i, j*. Under this condition, every individual exhibits the phenotype *x*_*i*_(*τ*) = 1*/α*, showing poor functionality as either sex, with both *W*_*m*_ and *W*_*f*_ ≪ 1. The population then evolves under mutation, selection, and recombination until clear sexual differentiation emerges or the number of generations exceeds 5×10^6^. We define that the two sexes are established when max(*W*_*f*_, *W*_*m*_) > 0.95 is satisfied for all individuals.

### Detection of sex-determining locus by association analysis

To identify sex-determining loci, we calculated the correlation between genotypes at each locus and the sexual phenotype of individuals. Because the phenotype in our model is an *n*-dimensional trait, we first summarized the degree of maleness of each individual by a single scalar score, *z*:

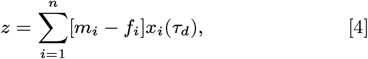

where *m*_*i*_ and *f*_*i*_ are the optimal expression levels of the *i*th gene for male and female functions, respectively. This score was evaluated based on the phenotype at the end of development. Individuals with more positive *z* values are considered more male-like, whereas those with more negative *z* values are more female-like. For each locus, we then calculated Kendall’s *τ*_*b*_ rank correlation between the mean genotypic value of each individual and its corresponding maleness score. As shown in Fig. 2d, one locus typically exhibits a particularly strong correlation, except during turnover periods, and we regarded that locus as the sex-determining locus.

### Detection of turnover events

To quantify the number of turnover events, we considered a turnover to be complete when the following three conditions were satisfied: (i) two sexes have been established in a population, (ii) alleles that show sex-biased distribution exist at the new sex-determining locus, and (iii) an allele is fixed at the previous sex-determining locus. For the second criterion, we regarded a focal allele as sex-biased when its allele count exceeded 400 in one sex and was zero in the other. Although these thresholds are somewhat arbitrary, we confirmed that automated detection based on them was largely consistent with manual inspection of each simulation run.

### Model for the scenario of degenerated sex chromosome

To investigate the evolution following the degeneration of Y or W chromosomes, we conducted simulations. For 1,000 replicate populations that successfully evolved sexual dimorphism within 5×10^6^ generations, we ran an additional 5×10^5^ generations to stabilize the sex-determining system. We then introduced a strongly deleterious mutation (selection coefficient *s*_*d*_ = −0.9) on all Y or W chromosomes in the population. This mutation was assumed to be completely linked to the ancestral sex-determining locus and to reduce individual viability. To incorporate selection against this mutation, we implemented an additional viability selection step that occurred after development but before mating. During this step, heterozygous individuals carrying this mutation die with probability |*s*_*d*_|, while homozygous carriers were lethal. Because the mutation was introduced only on Y or W chromosomes, it caused a strong bias in the sex ratio by reducing the frequency of the heterogametic sex, thereby strongly promoting turnover of the sex-determining system.

After the introduction of the deleterious mutation, simulations were continued for another 5×10^6^ generations to examine the potential for recovery from this perturbed state. Although we used this treatment of instantaneous degeneration to demonstrate our point, rather than the gradual degeneration typically observed in nature, it would provide valuable insight into the mechanisms enabling rapid fitness recovery.

## Data Availability

Codes used for simulations are available at https://github.com/TSakamoto-evo/sex_GRN.

---

**Table S1:** Variables and parameters in our model.

| variable | explanation | default | other values investigated |
| --- | --- | --- | --- |
| $t$ | the number of generations since the simulation starts | — | — |
| $\tau$ | developmental step | — | — |
| $x(\tau)$ | the expression of $n$ genes at $\tau$ | — | — |
| $N$ | population size (diploid) | 1,000 | — |
| $\alpha$ | the decay rate of gene expression | 0.2 | — |
| $\tau_m$ | the developmental step when the expressions start to affect fitness | 21 | — |
| $\tau_d$ | the developmental step when the expressions end to affect fitness | 40 | — |
| $\sigma$ | inverse of the strength of selection | 5.0 | strong selection: 3.0<br>weak selection: 7.0 |
| $\mu_g$ | mutation rate per locus for the type-G locus | $10^{-6}$ | low: $2 \times 10^{-7}$<br>high: $5 \times 10^{-6}$ |
| $\mu_b$ | mutation rate per locus for the type-B locus | $10^{-6}$ | low: $2 \times 10^{-7}$<br>high: $5 \times 10^{-6}$ |
| $\gamma_g$ | maximum effect size of a new mutation for the type-G locus | 0.1 | — |
| $\gamma_b$ | maximum effect size of a new mutation for the type-B locus | 0.1 | — |
| $\vec{M}, \vec{F}$ | optimal expression pattern in male and female, respectively | $\vec{M} = (10, 10, 0, 0, 10, 10, 0, 0)$<br>$\vec{F} = (10, 10, 0, 0, 0, 0, 10, 10)$ | divergent: $\vec{M} = (10, 10, 10, 10, 0, 0, 0, 0)$<br>$\vec{F} = (0, 0, 0, 0, 10, 10, 10, 10)$ |

**Fig. S1:**
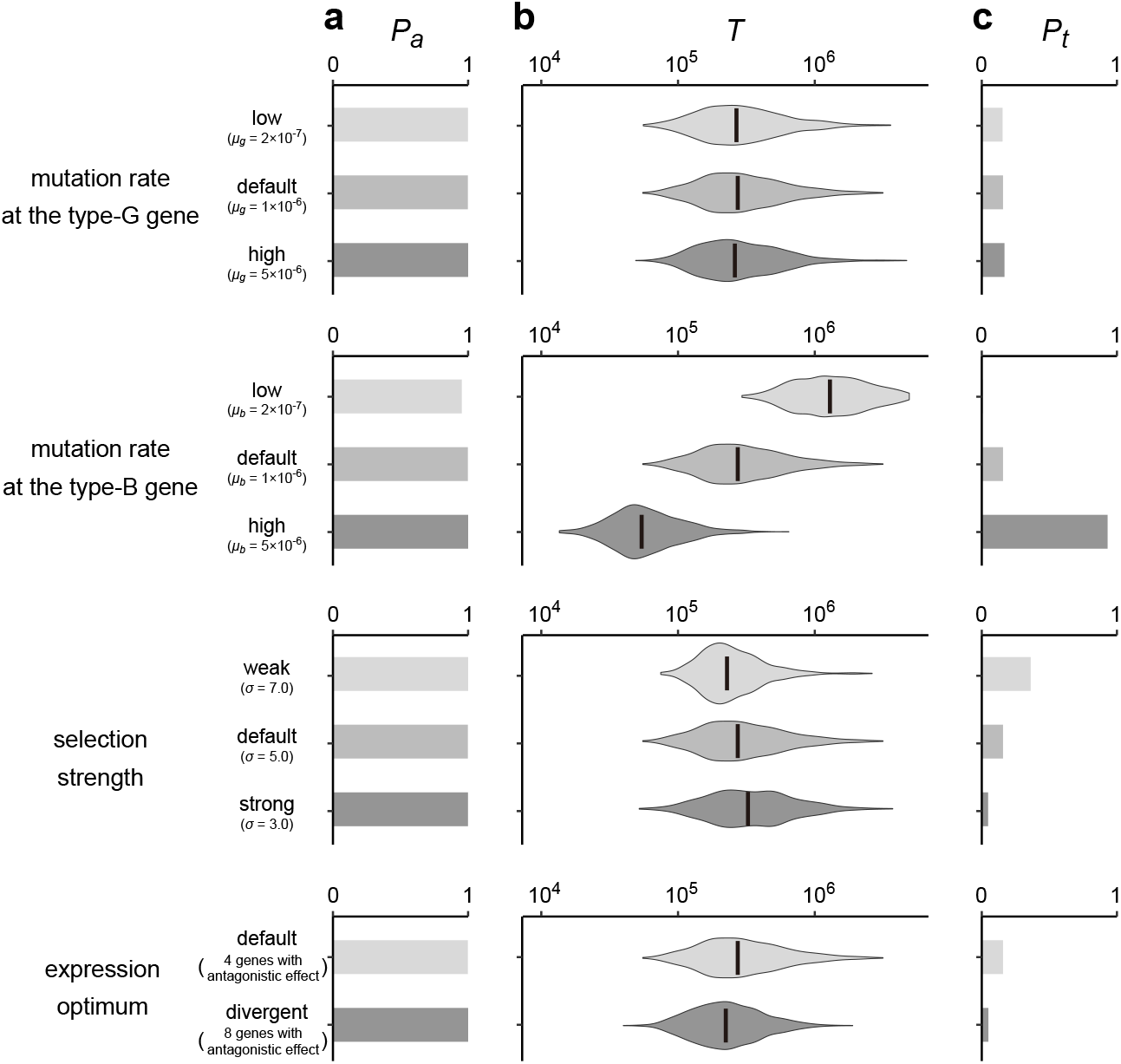
Evolution of anisogamy with various parameters. The effects of the mutation rate at the type-G locus, the mutation rate at the type-B locus, the selection strength and the expression optimum are investigated. (a) The proportion of runs in which the two sexes successfully evolved within 5 × 10^6^ generations, *P*_*a*_. (b) The distribution of the waiting time until the emergence of the two sexes, *T*. The vertical lines show the median of *T*. (c) The proportion of runs in which turnovers occur within 5 × 10^6^ generations after the emergence of the two sexes, *P*_*t*_. Only runs with successful anisogamy evolution were used for calculation. See also Table S1 for the examined parameters.

**Fig. S2:**
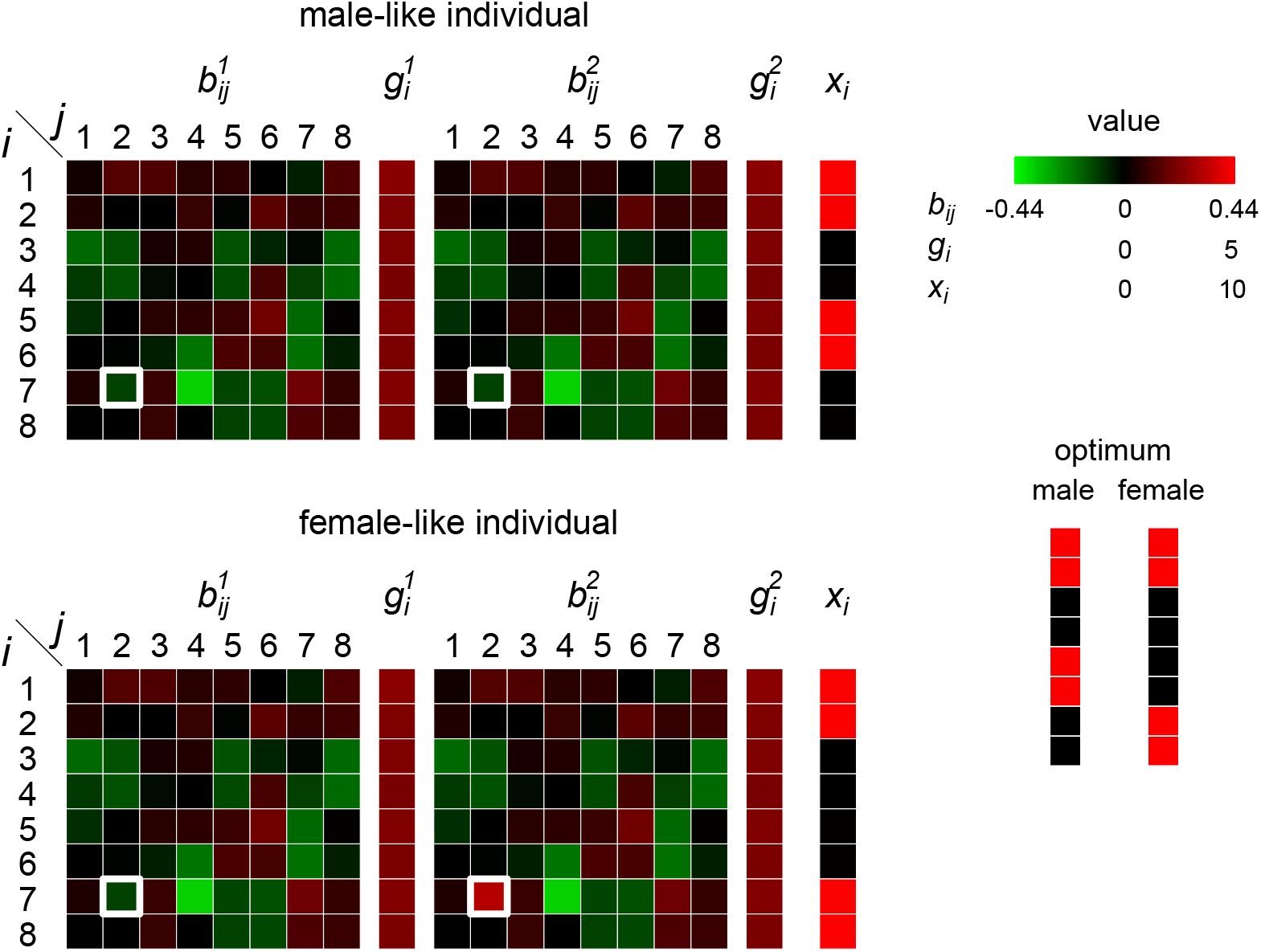
Genotypes after the evolution of anisogamy. Genotypes of representative male-like and female-like individuals at *t* = 2,000,000 are shown. The simulation run is the same as that in Fig. 2. The sex-determining locus (*b*_72_), highlighted by bold white rectangles, is the only notable difference between males and females.

**Fig. S3:**
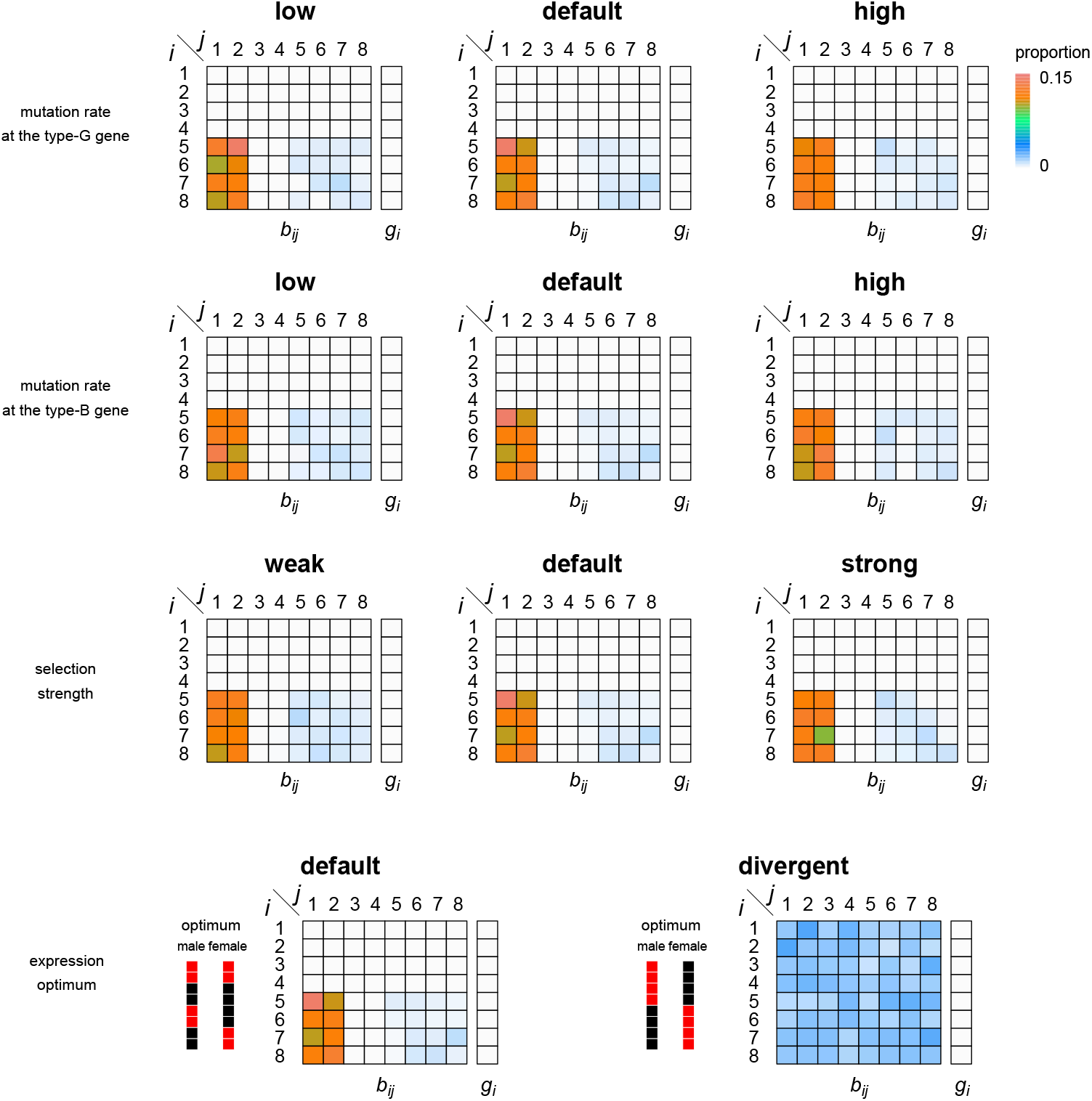
Frequency distribution of sex-determining locus in each parameter set. For examined parameter values, see Table S1.

**Fig. S4:**
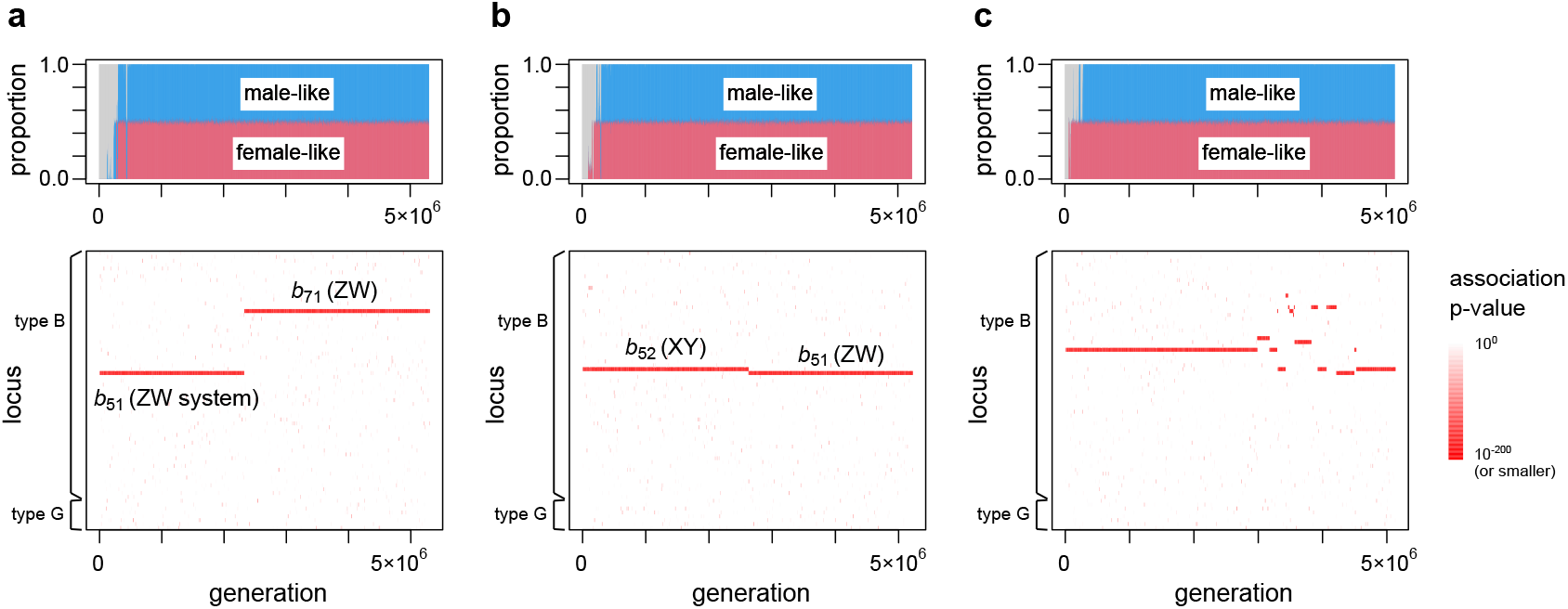
Examples of simulation runs showing different types of turnover events. (a) turnover without changing the heterogametic sex (ZW maintained), (b) turnover with XY-ZW switch, and (c) multiple turnovers in short time intervals. The color scale indicates the association p-value for each locus.

**Fig. S5:**
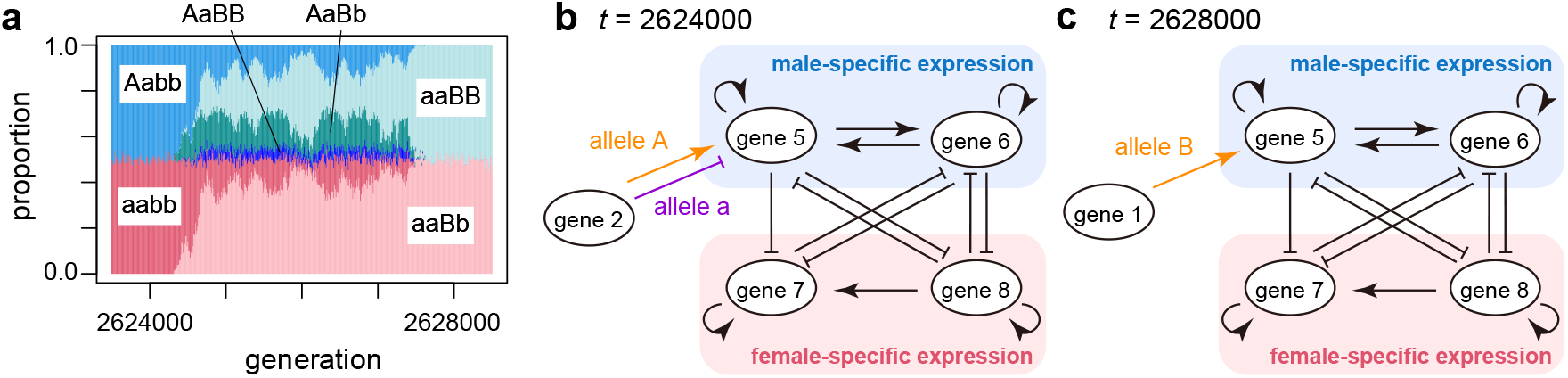
An example of the turnover with a transition from ZW to XY system. (a) The relative proportion of six major genotypes (Aabb, AaBB, AaBb, aaBB, aabb and aaBb) during the turnover. Other genotypes are very rare and colored in gray. The simulation run is identical to the one shown in Fig. S4b. (b, c) Gene regulatory relationships involved in the sexual differentiation before (b) and after the turnover (c). Gene regulations with |*b*_*ij*_ | > 0.05 are shown for *i, j* ∈ {5, 6, 7, 8}.

**Fig. S6:**
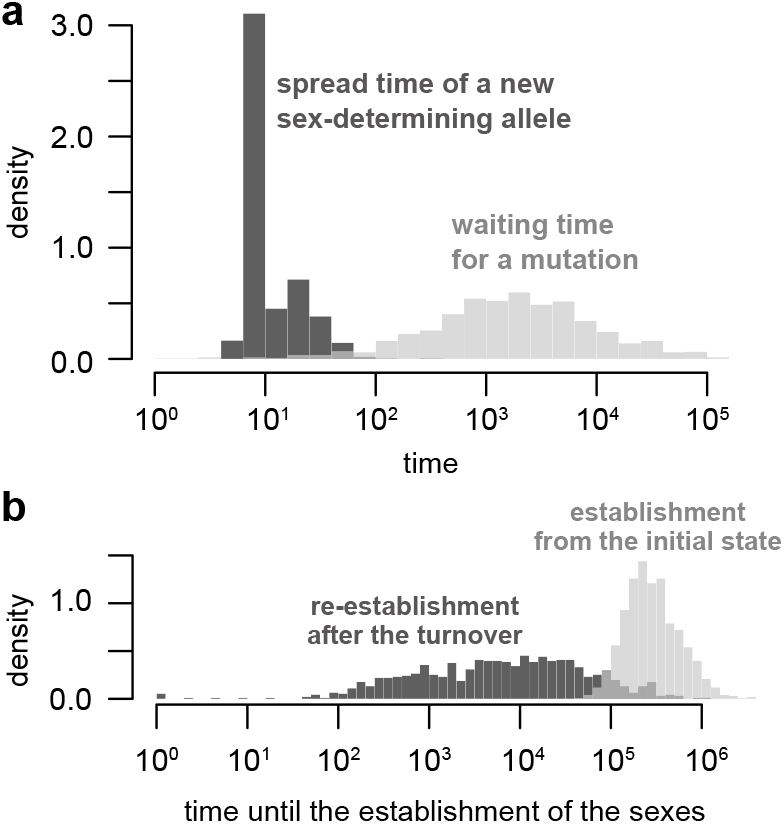
Detailed dynamics of the recovery process under the default parameter set. (a) Decomposition of the total time until the recovery completion. We recorded the time at which a new sex-determining allele originated (*t*_ori_). The time when the deleterious allele went extinct was denoted as *t*_purge_, while *t*_intro_ represents the time when the deleterious allele was introduced into the population (i.e., *t*_intro_ = *T* + 500,000). We then calculated the spread time of the new sex-determining allele (dark gray) as *t*_purge_ − *t*_ori_, and the waiting time for its origin (light gray) as *t*_ori_ − *t*_intro_. Seven cases were excluded from this analysis because the new sex-determining allele already existed at *t* = *t*_intro_ as standing variation. (b) Distribution of the number of generations until well-developed sexes are re-established (i.e.,, max(*W*_*m*_, *W*_*f*_) > 0.95) after the removal of the deleterious allele. Since this condition is already satisfied at *t* = *t*_purged_ in 95 cases, the results of the remaining 905 cases are shown here. For a comparison, the time until establishment from the isogametic state (*T*) is shown in light gray.

**Fig. S7:**
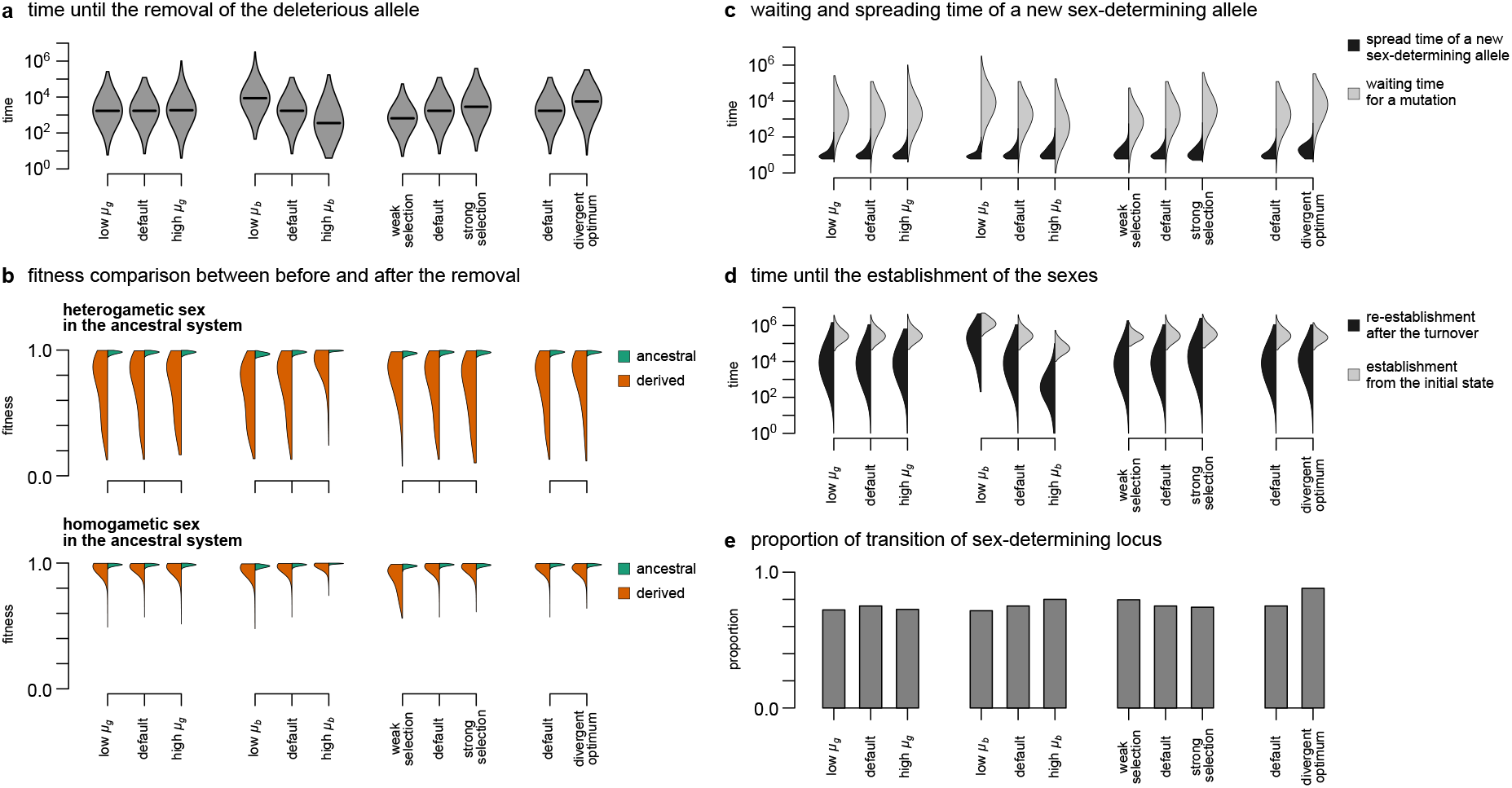
Effect of the parameters on the recovery dynamics from the degenerated state. (a) Distribution of the number of generations required to remove the deleterious allele. The data are from 1,000 simulation replicates where the two sexes are established within 5 × 10^6^ generations. The lines show the median. (b) Comparison of the fitness just before and after the removal of the deleterious allele. The mean fitness of the ancestrally heterogametic and homogametic sexes are shown in each panel. (c) Decomposition of the total time until recovery completion. The spread time is defined as the period between the origin of the new sex-determining allele and the extinction of the old sex-determining allele linked to the deleterious allele. The waiting time denotes the period between the introduction of the deleterious allele and the origin of the new sex-determining allele. See also the legend of Fig. S6. Cases in which the new sex-determining allele already existed at the time of deterioration as standing variation were excluded from this analysis. (d) Distribution of the number of generations until well-developed sexes are re-established (i.e.,, max(*W*_*m*_, *W*_*f*_) > 0.95) after the removal of the deleterious allele. We excluded the cases from the analysis in which this condition is already satisfied at the time of the removal of the deleterious allele. For a comparison, the time until establishment from the isogametic state (*T*) is shown in pale gray. (e) Proportion of the simulation runs in which the sex-determining locus underwent the turnover through the removal of the deleterious allele. See Table S1 for examined parameter values.

